# Ancestral ecological regime shapes reaction to food limitation in the Least Killifish, *Heterandria formosa*

**DOI:** 10.1101/2020.11.03.367128

**Authors:** Anja Felmy, Jeff Leips, Joseph Travis

## Abstract

In populations with contrasting densities of conspecifics, we often see genetically-based differences in life histories. The divergent life histories could be driven by several distinct agents of selection, including, amongst other factors, variation in per-capita food levels, the intensity of crowding-induced stress, rates of pathogen transmission, mate encounter rates, and the rates with which waste products accumulate. Understanding which selective agents act in a particular population is important as the type of agents can affect both population dynamics and evolutionary responses to density-dependent selection. Here we used a full-factorial laboratory experiment to examine whether two populations of a small live-bearing freshwater fish, characterised by high-density/low-predation or low-density/high-predation conditions, are adapted to different per-capita food levels. As expected, fish from the higher density regime handled food limitation better than those from the lower density regime. Although the lower food level resulted in slower growth, smaller body size, delayed maturation and reduced survival in both populations, especially survival to maturity showed a highly significant population x food-level interaction. At low food, 75% of fish from the low-density population died, compared to only 15% of fish from the high-density population. This difference was much smaller at high food (15% vs. 0% mortality), and was mediated, at least partly, through a larger size at birth of fish from the high-density regime. While we cannot preclude other agents of selection from operating differently in the study populations, we demonstrate that selection at higher density confers a greater ability to cope with low per-capita food availability.

## Introduction

Ecologists have long been interested in whether animal populations experiencing different regimes of population density will evolve adaptive differences in response to these conditions (Berec et al. 2018; Boyce 1984; Engen and Saether 2017; Macarthur and Wilson 1967; Mueller 1997; Pianka 1970; Wright et al. 2019). Careful experimental studies of laboratory populations have shown this to be the case (Bull et al. 2006; Mueller 1988). While associations between trait values and density regimes are well-documented in natural populations, compelling tests of whether such associations emerge from density-dependent selection are much rarer (Reznick et al. 2019; Travis et al. 2013).

One of the challenges in understanding the incidence and importance of density-dependent evolution is identifying the agents of selection behind adaptation to different population densities. The contrast between lower and higher density can include a host of ecological differences, depending upon the specific case. Higher density can mean, among other factors, lower per-capita food levels, a higher rate of stressful social interactions, higher rates of pathogen transmission, greater competition for mates, and, in some systems, higher rates of accumulation of waste products (Berec et al. 2018; Than et al. 2020). Laboratory studies with *Drosophila* have shown that nearly all of these agents can be acting (Joshi et al. 1996; Joshi and Mueller 1988; Joshi and Mueller 1993; Joshi and Mueller 1996; Joshi et al. 1998; Mueller 1988; Mueller 1990; Sarangi et al. 2016).

Understanding how selection acts differently in different density regimes is not a mere exercise in detail. The responses to density-dependent selection and their consequences for subsequent population dynamics depend on the demographic details of selection, for example regarding the age classes primarily affected (Charlesworth 1994; Engen and Saether 2016; Mueller et al. 2000), and on which traits respond to which selection pressures (Engen et al. 2020; Tung et al. 2019).

Among the most obvious selection pressures that differ between density regimes is per-capita food availability. Laboratory studies have shown that populations selected to evolve at higher densities respond with adaptations in feeding rate (Joshi and Mueller 1996; Mueller 1990) and digestive efficiency (Sarangi et al. 2016), consistent with a response to reduced food availability. In natural populations of Trinidadian guppies (*Poecilia reticulata*), populations at chronically higher densities differ from those at chronically lower densities in having an expanded diet (Bassar et al. 2017; Zandona et al. 2011), reduced nitrogen excretion rates (El-Sabaawi et al. 2015), lower resting metabolic rates (Auer et al. 2018), and stoichiometric relations indicative of resource limitation (El-Sabaawi et al. 2012). These differences correspond with what one might expect as responses to relative food scarcity (higher density) or abundance (lower density). Experiments with Hart’s killifish (*Rivulus hartii*) have shown that populations living at consistently higher conspecific densities are more capable of growth and development at low food levels than populations at consistently lower densities (Walsh and Reznick 2010).

Despite these examples, it cannot simply be assumed that adaptation to lower per-capita food levels will be evident in all natural populations experiencing higher densities. For one reason, the social environment in a high-density population can affect vital rates independently of food levels (Gutierrez et al. 2020; Leatherbury and Travis 2019) and can be a powerful selective agent in its own right. For another, populations living at different densities need not be experiencing different per-capita food levels if primary productivity varies among locations, making the ratio of food supply to metabolic demand, and not food level per se, the important ecological variable (Wilbur 1977). Finally, the detrimental effects of higher density can depend upon abiotic factors that affect the metabolic demand of an organism (Travis and Trexler 1986; Warner et al. 1991), with the result that the mere observation of consistent differences in density need not imply consistent differences in per-capita resource availability. Only experiments can assess whether adaptation to lower per-capita food levels is an outcome of natural selection operating at high population densities.

The Least Killifish, *Heterandria formosa,* offers an excellent opportunity to investigate this issue. This fish is found throughout the lower coastal plain of the southeastern United States. Populations in north Florida occur in a wide variety of habitats, from freshwater springs to lakes, ponds, and swamps in river bottoms (MacRae and Travis 2014). In all habitats, *H. formosa* occupies the shallow littoral zone and is a primary consumer (Aresco et al. 2015). Adults are small, with male standard length (SL, the distance from the tip of the snout to the hypural plate in the tail) varying, typically, from 9 mm to 15 mm and female SL varying from 10 to 25 mm. Reproduction is placental; females provide almost all nourishment for their embryos after fertilisation.

Prior work has shown that population density is dramatically higher and per-capita predation risk lower at Wacissa River (WR) than Trout Pond (TP; Leips and Travis 1999; MacRae and Travis 2014; Richardson et al. 2006). Size at maturity is smaller, fecundity is higher, and average offspring size lower in TP, distinctions based in genetic differences (Leips et al. 2000). The different life histories appear to be adaptations to the different regimes of density and predation risk that each population experiences, with higher predation and lower density favouring earlier maturation, greater fecundity and smaller individual offspring. While there are many abiotic differences between the two locations, there is no evidence that the two populations differ in their responses to them (Hale and Travis 2015), reinforcing the conclusion that the different life histories are adaptations to different biotic environments (Leips et al. 2000; Schrader and Travis 2012).

Mescocosm experiments implicate one or more factors associated with the two populations’ characteristic differences in population density as major agents of selection. Experiments manipulating population density (Leips et al. 2009; Leips et al. 2013) and genetic composition (Leips et al. 2000) showed that the two populations differed in their response to density, with the reproductive traits of individuals from the Wacissa River, the low-predation/high-density population, being less sensitive to the depressant effects of density and the degree of sensitivity being proportional to the initial dosage of Wacissa River alleles (Leips et al. 2000).

It is unclear if the reduced sensitivity to higher density in the WR fish reflects an adaptation to lower per-capita food levels. Trout Pond is the more productive habitat (Aresco and Travis, unpublished data), suggesting that the higher density of WR fish, which is a function of lower predation risk (MacRae and Travis 2014; Richardson et al. 2006), does create lower per-capita food levels. However, laboratory experiments indicate that social stress is at least as strong an effect on female reproduction rate as reduced food levels (Leatherbury and Travis 2019). In this paper we report the results of testing whether adaptation to different per-capita food levels plays a role in how these two populations have adapted to different density regimes.

## Materials and Methods

### Experimental design

We collected adults from Trout Pond, Wakulla County, Florida, and from Wacissa River, Jefferson County, Florida, in September 1993 and used them to establish breeding colonies in the laboratory. We housed the adults from each population in two 76-litre aquaria per population and collected F1 offspring as they were born. Those offspring were raised in 38-litre aquaria until maturity, when we removed females and mated them to males from different aquaria to minimise inbreeding. We isolated pairs in small aquaria and collected their offspring at birth for the experiment. We began the experiment in February 1994.

The experiment was a factorial design in which we raised offspring to maturity from either population (TP or WR) at one of two food levels, designated “high” or “low.” We set the initial daily “high” food ration as 4 mg and “low” as 1 mg. We increased the food ration as individuals grew to a maximum by day 63 of 20 mg/day at “high” and 5 mg/day at “low.” “Low” food was always set as one-quarter of “high.” We considered a male to be mature when his gonopodium (modified anal fin used as an intromittent organ) was fully formed; we considered a female to be mature when the characteristic black spot appeared on her anal fin. Sample sizes are provided in Table 1.

**Table 1.** Number of focal individuals as a function of population of origin, experimental food level, survival to sexual maturity, and sex.

| Population | Treatment | Initial<br>sample size | No. Survivors |  | No. Deaths |
| --- | --- | --- | --- | --- | --- |
|  |  |  | Male | Female | Sex unknown |
| Trout Pond | High food | 20 | 9 | 8 | 3 |
|  | Low food | 20 | 0 | 5 | 15 |
| Wacissa River | High food | 20 | 12 | 8 | 0 |
|  | Low food | 20 | 10 | 7 | 3 |

Only fish that survived could be sexed.

### Measurement of life-history traits

We measured each individual fish at birth, at 14 days, 28 days, 42 days, and at sexual maturity. We used standard length (SL) as a measure of body size. To measure each individual’s SL, we removed the fish from its aquarium, placed it in a small petri dish against a standard grid, photographed it, and returned it to its aquarium. No individual died as a result of this procedure. Standard length and dry mass were strongly correlated (Pearson’s product-moment correlation: *r* = 0.76, *t*_18_ = 5.01, *p* < 0.0001, n = 20 fry measured at birth). Photographic length measurements were highly repeatable (repeatability *r* = 0.99, effect of individual identity in a simple ANOVA: *F*_36,37_ = 381.3, *p* < 0.0001, *n* = 37 individuals that were measured twice).

### Statistical analysis

Size at birth was approximately normally distributed, as judged from a quantile-quantile plot (QQ-plot) and a two-sided Kolmogorov-Smirnov test (*D* = 0.10, *p* = 0.42). We therefore analysed it using a generalised linear mixed model with Gaussian errors. As predictors we included the population of origin (Trout Pond vs. Wacissa River), the experimental food level (high vs. low), maternal size (continuous) and the interaction between population and maternal size. The sole random effect was maternal identity. It is not possible to determine the sex of a juvenile from external morphology and so the sex of fish that died before attaining sexual maturity remains unknown. Given the differential survival of fish from TP and WR, and of fish assigned to the high- and low-food treatment, sex as a categorical predictor with three levels (female, male, unknown) was confounded with both population identity and food level. We therefore tested for a size difference between males and females in a separate model using only the individuals of known sex. All remaining predictors in this model were as before. As the QQ-plot identified four data points as potential outliers (size at birth > 8.0 mm), we fitted additional models after excluding these data points.

Survival to sexual maturity was analysed using a generalised linear mixed model with binomial errors. As fixed effects we included the population of origin, the experimental food level, size at birth (continuous), and maternal size (continuous), and as random effects, maternal identity. We were unable to include the interaction between population identity and food level, because the main effects of population (92.5% surviving fish from WR vs. 55.5% from TP) and of the food treatment (92.5% surviving fish at high food vs. 55.5% at low food) were exactly the same, causing the model to suffer from computational problems. We therefore used a Chi-square test, which proved robust to the mirror-image effects of population and food level, to compare the number of dead and surviving fish among the four experimental groups. To identify the cells in the contingency table that accounted for most of the difference between expected and observed values, we computed the relative contribution of each cell to the total Chi-square score as 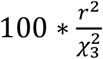, where *r* was the Pearson residual and 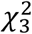 the Chi-square score.

To analyse size during adolescence and at sexual maturity, we performed two analyses. The first one was a generalised linear mixed model with size at 14 days as the dependent variable, which included all individuals that survived to and were measured at that age (n = 66). Size at 14 days was approximately normally distributed (QQ-plot; *D* = 0.14, *p* = 0.17), so models with Gaussian errors were fitted. For predictors, we used the population identity, food level, their interaction, maternal size, and size at birth, plus maternal identity as a random effect. Originally, we included interactions between food level and size at birth, between population and size at birth, and between population and maternal size; we dropped these effects from the final model because they had no impact on the response (elimination criterion: ratio of *χ*^2^ over its degrees of freedom < 1 in log-likelihood ratio tests comparing a model with and without the interaction in question). We could not include sex as a predictor due to a small but non-negligible number of fish with unknown sex (n = 8). Again, the five fish that were largest at 14 days (> 9 mm) appeared to be outliers, based on a QQ-plot. To explore their influence on results, we also fitted a model without them.

We additionally analysed the three juvenile sizes and size at maturity with a repeated measures analysis of variance, using function “aov” in R (R Core Team 2020). We included only individuals that survived to maturity and were measured on all four occasions (n = 55). All these individuals were of known sex. The predictors were population, food level, age category, sex, size at birth, maternal size, all the two-way interactions between age and these predictors (except the age x maternal size interaction), and maternal identity. Our formula contained individual identity as the single error term, to account for repeated measures of fish. The error term specified two error strata, with appropriate models fitted within each stratum. We tested the effects of age and all its interactions within individuals, while we tested all the other effects between individuals. The low number of survivors that were from Trout Pond, kept at the lower food level, and had no missing data (n = 3) meant that we were unable to fit the interaction between food level and population. We also initially fitted, and then dropped from the final model due to non-significance, the interactions between age and maternal size, between population and size at birth, between food level and sex, and between population and sex (elimination criterion: *F*-value < 1).

The age at sexual maturity was approximately normally distributed (QQ-plot; *D* = 0.11, *p* = 0.52), and so we analysed this variable with generalised linear mixed models with Gaussian errors. Our final model included the population, food level, size at birth, sex, and the interaction between food level and size at birth as fixed effects, and maternal identity as a random effect. We did not fit more predictors to ensure sufficient statistical power given the reduced sample size in this analysis (n = 57 fish with known age at maturity). We therefore did not include non-experimental (i.e. purely correlative) predictors in our model that, upon visual inspection, appeared unrelated to the age at sexual maturity (e.g. maternal size, size at maturity, and the interaction between size at birth and population). In addition, we could not fit the interactions between population and food level, population and sex, and food level and sex, because in each of these cases one or several treatment levels had insufficient sample sizes (i.e. < 10 data points). Three data points (age at maturity < 36 days and > 85 days) showed slight deviations from normality on the QQ-plot, and we fitted an additional model after excluding them.

We assessed the fit of all models that converged with diagnostic plots. The reference levels for categorical predictors were Trout Pond, high food, female. In models that included maternal identity as a random effect, we obtained a *p*-value for maternal identity with log-likelihood ratio tests comparing the full model to one without random effects.

We performed all statistical analyses in R 4.0.0 (R Core Team 2020). Unless stated differently, we fitted generalised linear mixed models using function “glmmTMB” in R-package “glmmTMB” (Brooks et al. 2017). Values are given as mean ± SD. Binomial standard errors for survival rates were computed according to Zar (1996) as 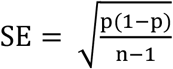, where p was the proportion of fish that survived and n the total number of fish. Scatter plots were prepared using R-package “beeswarm” (Eklund 2016) and depict individual data points superimposed on boxplots. White triangles on boxplots show group means.

## Results

### Size at birth

Fish from Wacissa River were about 15% larger at birth than fish from Trout Pond (*b* = 4.04, *z* = 2.25, *p* = 0.0245, Fig. 1a). They were also slightly larger when assigned to the high-rather than the low-food treatment, although this effect is coincidental, as fish were measured before they were fed for the first time (*b* = −0.36, *z* = −3.08, *p* = 0.0021, Fig. 1b). There was no overall effect of maternal size on offspring size at birth (*b* = 0.09, *z* = 1.44, *p* = 0.15), but in fish from WR larger offspring tended to be born to smaller mothers (interaction between maternal size and population: *b* = −0.14, *z* = −1.78, *p* = 0.07, Fig. 1c). Maternal identity, included as a random effect, explained a significant amount of variation in offspring size at birth (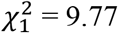, *p* = 0.0018). We tested for a size difference between the sexes using only the subset of fish with known sex, and found that males and females did not differ in size at birth (*b* = −0.25, *z* = −1.62, *p* = 0.11, Fig. 1d). The inclusion of sex as a predictor left the remaining model results largely unchanged (data not shown).

**Figure 1.**
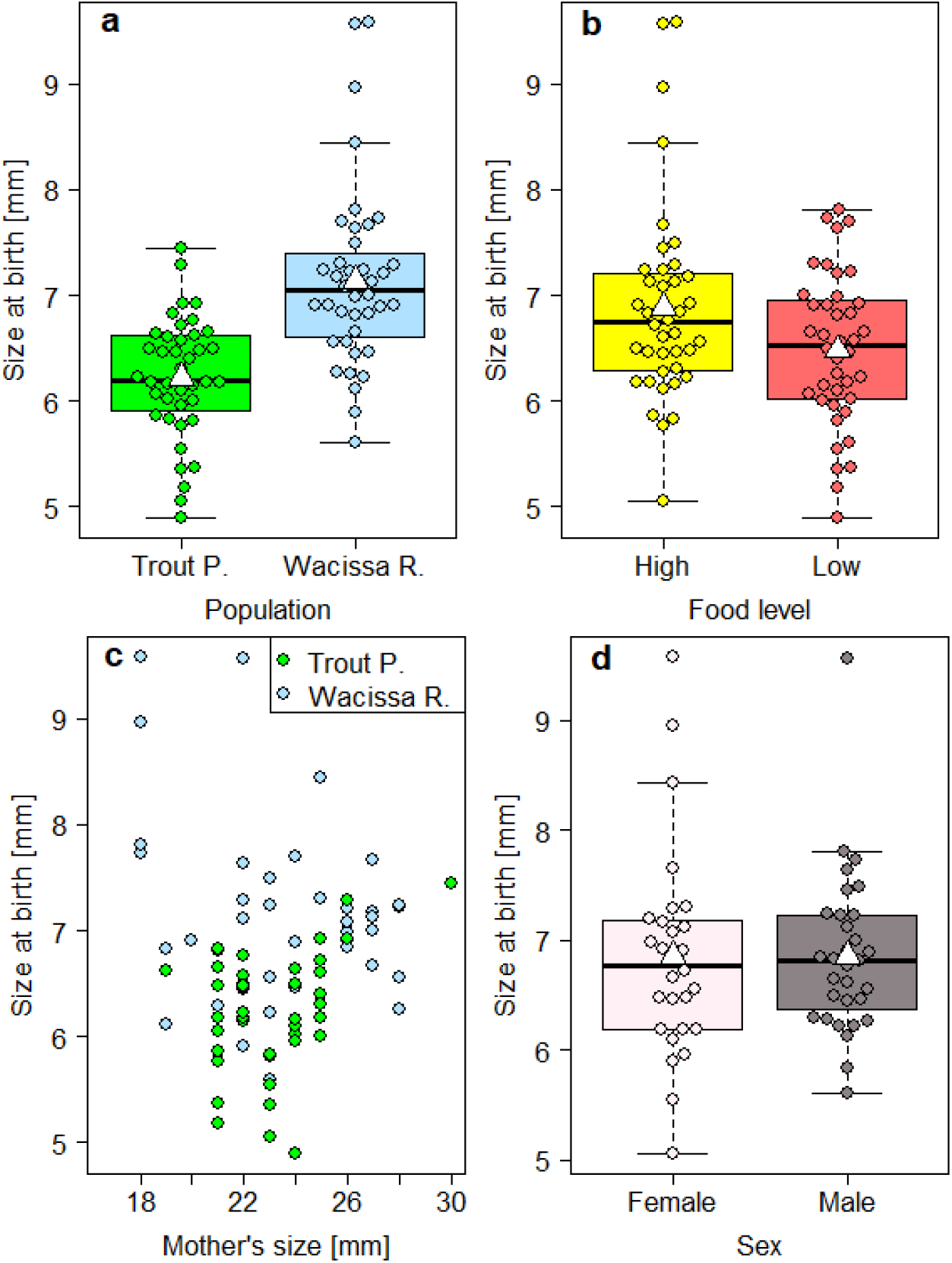
Size at birth was affected by population of origin, and weakly associated with maternal size. The effect of food levels is coincidental and disappeared after the four largest individuals (> 8mm) were excluded. Males and females did not differ in size at birth. In (d), fish of unknown sex are excluded.

Some of these results were contingent on four exceptionally large fry (> 8.0 mm), which were identified as potential outliers in a QQ-plot. When we excluded these fry, the population of origin still had a strong effect (*b* = 2.94, *z* = 2.23, *p* = 0.0258, Fig. 1a), but the coincidental effect of food level disappeared (*b* = −0.17, *z* = −1.53, *p* = 0.13, Fig. 1b). Moreover, the main effect of maternal size became significant, indicating that larger mothers were likely to produce larger offspring (*b* = 0.10, *z* = 2.12, *p* = 0.0340, Fig. 1c). This relationship was evident in fish from TP; in fish from WR, there was still a slight tendency for larger mothers to have smaller offspring (interaction between maternal size and population: *b* = −0.10, *z* = −1.73, *p* = 0.08). When using the outlier-free dataset, maternal identity did not predict size at birth (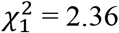, *p* = 0.12). The size difference between males and females remained non-significant (*b* = −0.02, *z* = −0.16, *p* = 0.88, Fig. 1d).

### Survival to maturity

Twenty-six percent of fish died before reaching sexual maturity. The mortality rate was highest in the two weeks after birth (16.3%), intermediate between 14 and 28 days of age (7.5%), and lowest between 28 days and the age when fish attained maturity (2.5%). Survival to maturity was significantly higher among fish from WR (92.5%) than among fish from TP (55.0%, *b* = 2.95, *z* = 2.91, *p* = 0.0037, Fig. 2a). An effect of the same exact size was caused by the food treatment, with 92.5% surviving fish under high- and only 55.5% under low-food conditions (*b* = −2.94, *z* = −3.57, *p* = 0.0004, Fig. 2a). The association between size at birth and survival was non-significant (*b* = 0.17, *z* = 0.27, *p* = 0.79). However, once population identity was removed from the list of predictors, a larger size at birth was associated with increased survival (*b* =1.22, *z* =2.55, *p* = 0.0107, Fig. 2b), showing that the higher survival rate of fish from WR was, at least in part, mediated through a larger size at birth. In addition, survival tended to be slightly higher in fish born to larger mothers (*b* = 0.28, *z* = 1.67, *p* = 0.10, Fig. 2c). Maternal identity did not explain any variation in offspring survival (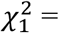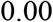, *p* = 1.00).

**Figure 2.**
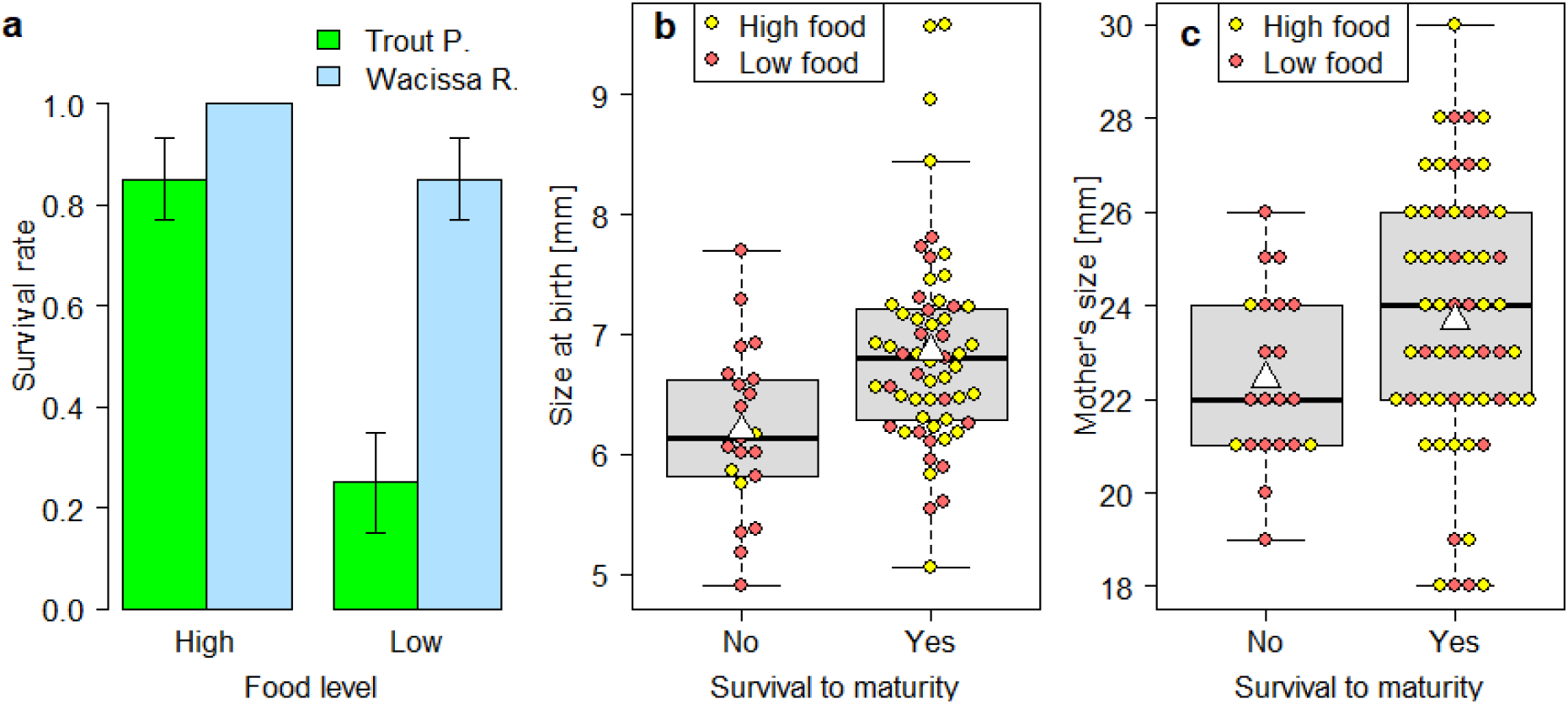
Survival was reduced at the lower food level, particularly in fish from Trout Pond, and positively associated with size at birth and potentially maternal size. In (a), sample size is 20 fish for each vertical bar, and error bars are binomial standard errors (± 1 SE). Note that survival was 100% for fish from Wacissa River kept at high food, and the binomial standard error consequently zero.

Fish from TP were more sensitive to food limitation than fish from WR: at the lower food level, 75% of fish from TP died, compared to only 15% of fish from WR (Fig. 2a). Mortality rates were much lower at the higher food level and not so different between populations: 15% in TP and 0% in WR. This population identity × food level interaction was evident in a Chi-square test (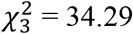, *p* < 0.0001), of which the residuals showed an excess of dead (Pearson residual *r* = 4.26) and concomitant lack of surviving fish (*r* = −2.54) that were from TP and kept at the lower food level, and a lack of dead fish that were from WR and kept at the higher food level (*r* = −2.29). Accordingly, the cells in the contingency table that contributed the most to the total Chi-square score were dead fish / TP / low-food (52.8%), surviving fish / TP / high-food (18.8%), and dead fish / WR / high-food (15.3%). Together, these cells accounted for 86.9% of the observed difference between expected and observed frequencies.

### Size during adolescence and at maturity

Fish from WR were about 15% larger than fish from TP at 14 days of age (*b* = 0.41, *z* = 2.13, *p* = 0.0330), and fish on the high food ration were about 14% larger at this age than fish on the low food ration (*b* = −0.43, *z* = −2.22, *p* = 0.0264, Fig. 3a). There was no interaction between population and food level (*b* = −0.30, *z* = −1.26, *p* = 0.21, Fig. 3a). At that young age, an individual’s size was still strongly correlated with its size at birth (*b* = 1.00, *z* = 10.75, *p* < 0.0001, Fig. 3b), and tended to be positively associated with maternal size (*b* = 0.05, *z* = 1.78, *p* = 0.08). The effect of maternal identity was non-significant (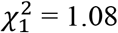, *p* = 0.30). Sex could not be included in the model, but a visual inspection of the data tentatively suggests that males and females did not differ in size at age 14 days, while fish that died as juveniles may potentially have been relatively small (Fig. 3c).

**Figure 3.**
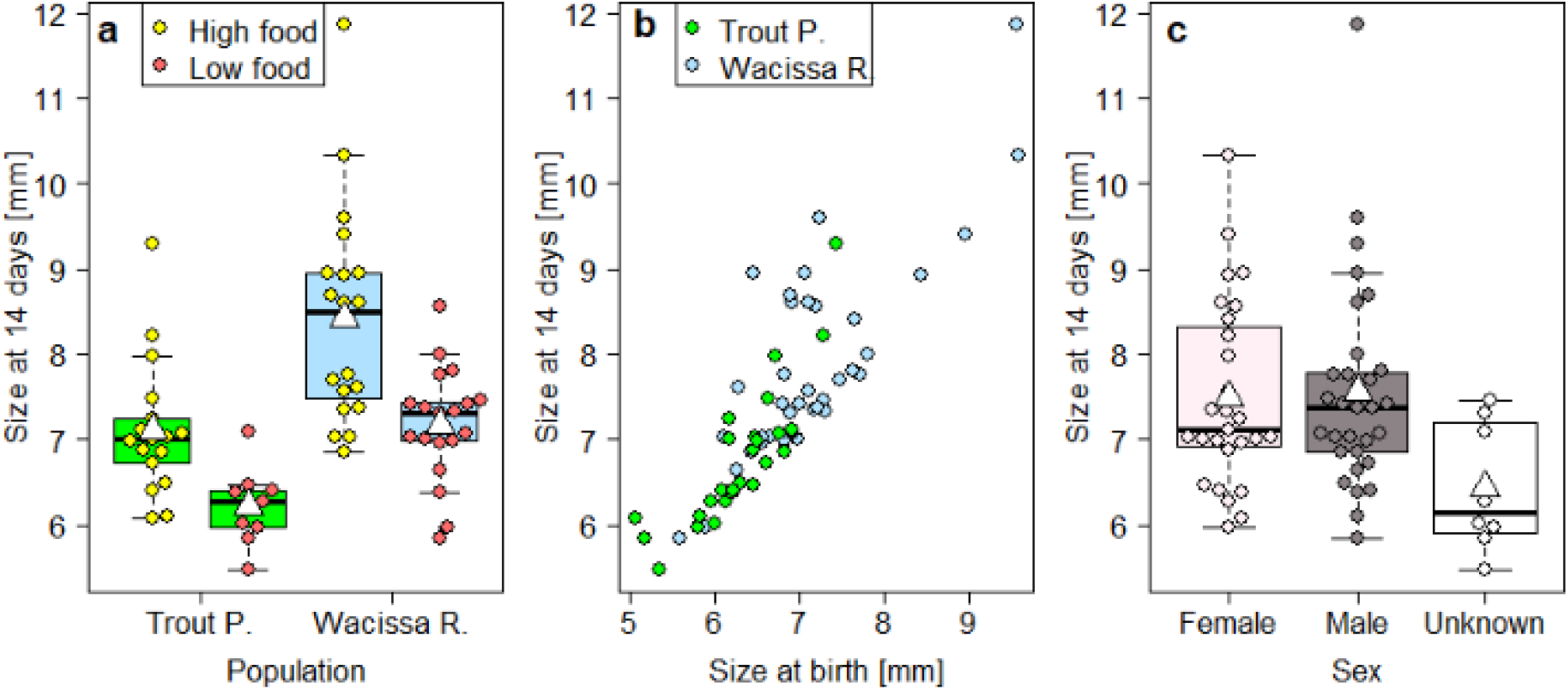
Size at 14 days was reduced at the lower food level and in fish from Trout Pond, and was highly correlated to size at birth. The relationship between size at 14 days and sex (c) is shown for illustrative purposes only, as the low number of fish that did not survive to sexual maturity and hence could not be sexed (n = 8) precluded its inclusion in the statistical model.

When excluding the five largest fish (> 9 mm), identified as potential outliers, the influence of food level grew stronger (*b* = −0.50, *z* = −3.46, *p* = 0.0005) and the effect of maternal identity became significant (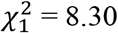, *p* = 0.0040). The correlation with maternal size disappeared altogether (*b* = 0.03, *z* = 0.84, *p* = 0.40). The other model results remained largely unchanged (population: *b* = 0.46, *z* = 2.52, *p* = 0.0116; population-food-level interaction: *b* = −0.09, *z* = −0.51, *p* = 0.61; size at birth: *b* = 0.75, *z* = 7.59, *p* < 0.0001).

A joint analysis of all juvenile sizes and size at sexual maturity (model results in Table 2) showed that fish from TP were always smaller (significant main effect of population), even at maturity, yet did not grow more slowly than fish from WR (non-significant age x population interaction, Fig. 4a), suggesting that population differences in juvenile size stemmed from the differential size at birth. Fish did grow more slowly under low-food conditions (significant main effect of food, and age × food interaction, Fig. 4b). The overall effect of sex was non-significant, owing to the fact that males and females did not visibly differ in size at 14, 28, and 42 days of age, but once they reached maturity males were considerably larger than females (significant age x sex interaction, Fig. 4c). Juvenile size also showed a positive statistical association with size at birth, which was most pronounced at 14 days and gradually disappeared at older ages, until no effect was left at sexual maturity (significant main effect of size at birth, and age x size-at-birth interaction, Fig. 4d). Neither maternal size nor maternal identity had a bearing on somatic growth (Table 2).

**Table 2.** Analysis of variance with repeated measures of juvenile sizes and size at sexual maturity.

|  | Df | Sum Sq | Mean Sq | *F*-value | *p*-value |
| --- | --- | --- | --- | --- | --- |
| Error: between individuals |  |  |  |  |  |
| Population | 1 | 38.61 | 38.61 | 11.87 | *0.0020* ** |
| Food level | 1 | 145.51 | 145.51 | 44.74 | *< 0.0001* *** |
| Sex | 1 | 2.58 | 2.58 | 0.79 | 0.38 |
| Size at birth | 1 | 43.64 | 43.64 | 13.42 | *0.0012* ** |
| Maternal size | 1 | 9.83 | 9.83 | 3.02 | 0.09 |
| Maternal identity | 24 | 86.24 | 3.59 | 1.11 | 0.40 |
| Residuals | 25 | 81.31 | 3.25 |  |  |
| Error: within individuals |  |  |  |  |  |
| Age | 3 | 927.3 | 324.11 | 324.08 | *< 0.0001* *** |
| Age x Population | 3 | 0.5 | 0.18 | 0.18 | 0.91 |
| Age x Food level | 3 | 16.7 | 5.56 | 5.56 | 0.0012 |
| Age x Sex | 3 | 62.4 | 20.79 | 20.78 | *< 0.0001* *** |
| Age x Size at birth | 3 | 19.6 | 6.55 | 6.55 | *0.0003* *** |
| Residuals | 150 | 150.0 | 1.00 |  |  |

The analysis includes standard lengths measured at 14 days, 28 days, 42 days, and at sexual maturity. Only individuals that survived to maturity and were measured on all four occasions were included (*n* = 55). Standard length was measured in mm. Italics: *p* < 0.05. Df: degrees of freedom, Sum Sq: sum of squares, Mean Sq: mean squares, SD: standard deviation.

**Figure 4.**
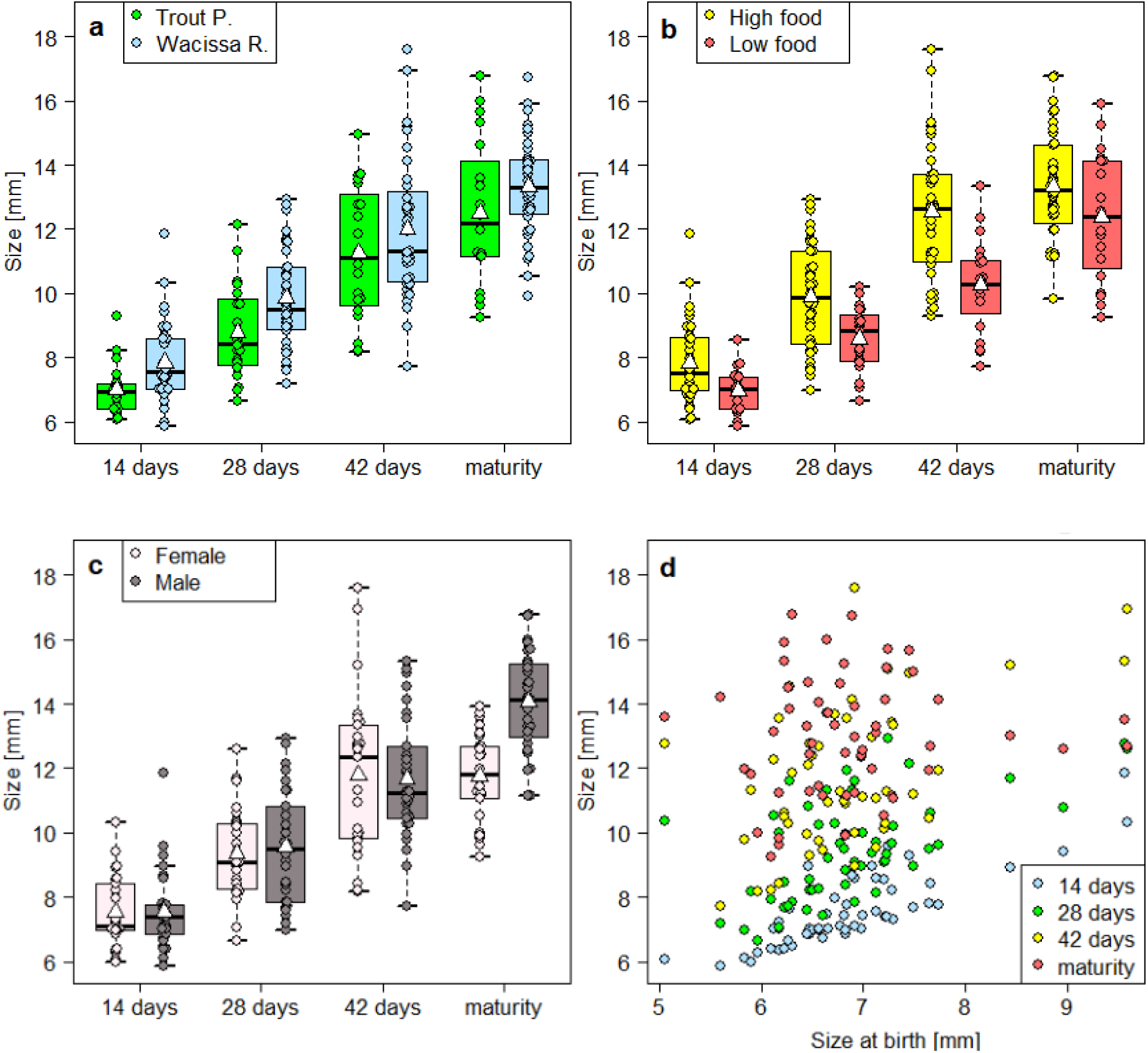
During adolescence and at maturity, body size was lower for fish from Trout Pond, for fish fed a low-quantity died, and, at maturity, for females. The effect of size at birth (d) gradually disappeared as fish grew older. This analysis only included fish that reached sexual maturity (n = 55).

### Age at sexual maturity

Fish from the two populations did not differ from each other in age at maturity (*b* = −2.77, *z* = −0.96, *p* = 0.34, Fig. 5a), but attained sexual maturity about 29% later when fed a low-quantity diet (*b* = 73.78, *z* = 3.10, *p* = 0.0019, Fig. 5b). The interaction between population and food level could not be analysed statistically due to the poor survival of fish from TP under low-food conditions (n = 4). On average, males matured substantially later than females (64 ± 13 days vs. 48 ± 13 days: *b* = 19.83, *z* = 8.21, *p* < 0.0001, Fig. 5c). Although there was no main effect of size at birth (*b* = −2.33, *z* = −1.34, *p* = 0.18), its interaction with the feeding regime was significant (*b* = −8.56, *z* = −2.43, *p* = 0.0153, Fig. 5d), indicating that fish kept at the lower food level reached sexual maturity earlier when they were born relatively large. Maternal identity did not explain any variation (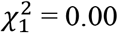, *p* = 1.00). The exclusion of two females that matured very early (29 and 32 days, respectively) and of a male that matured very late (90 days), which were identified as potentially influential data points in a QQ-plot, did not change these results (data not shown). The limited sample size (57 fish survived to sexual maturity) prevented us from fitting additional predictors, but a visual inspection of the data suggested that age at maturity was independent of maternal size (*r* = −0.04, *t*_55_ = −0.29, *p* = 0.77; Fig. 5e) and size at maturity (*r* = 0.18, *t*_55_ = 1.39, *p* = 0.17; Fig. 5f).

**Figure 5.**
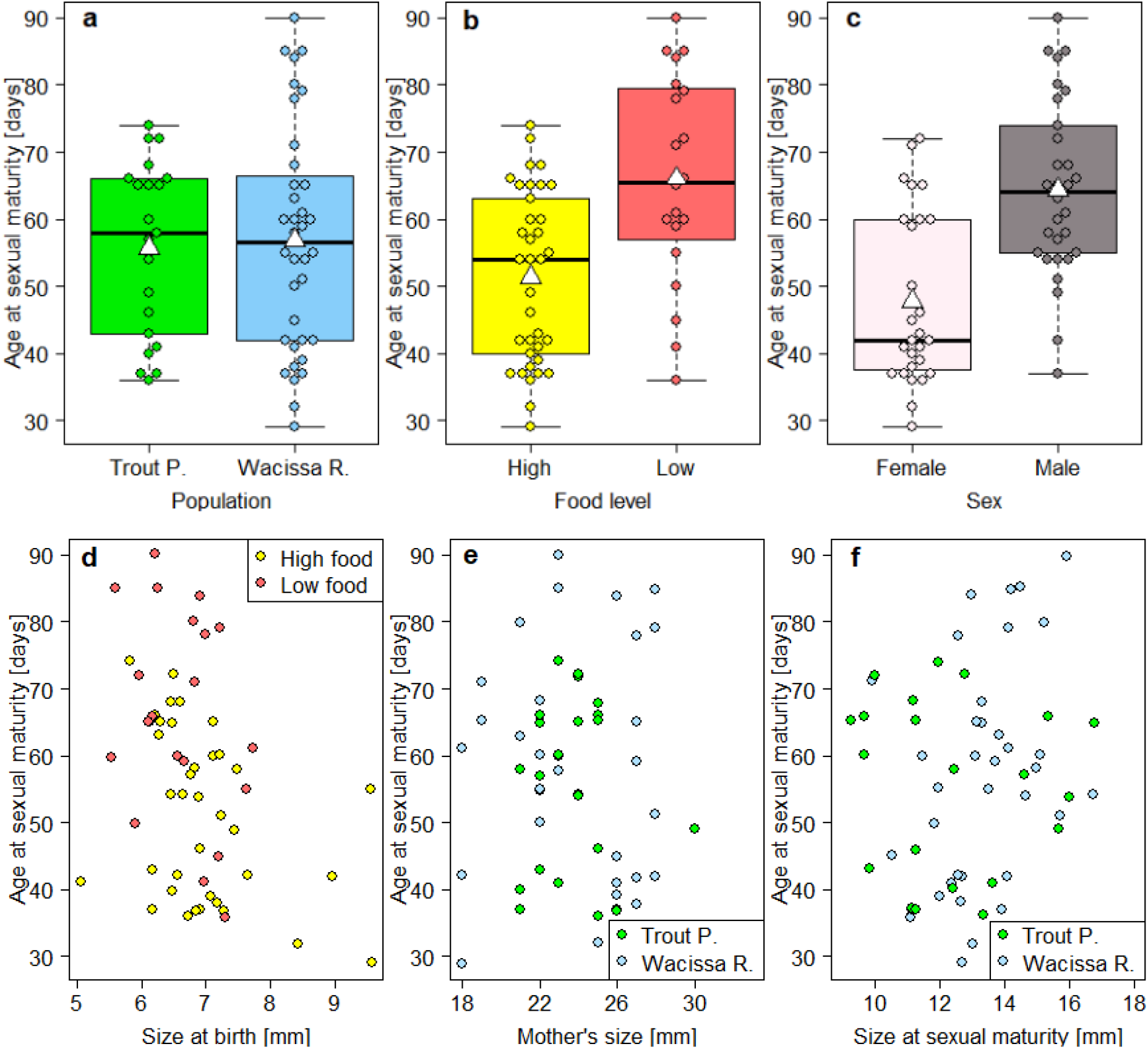
The age at sexual maturity was delayed at the lower food level, especially in fish that were small at birth, and was earlier in females than in males. There was no effect of population of origin. The relationships of age at maturity with maternal size (e) and with size at maturity (f) are shown for illustrative purposes only, and were not included in the statistical model.

## Discussion

Our results suggest that the divergent life histories of Least Killifish living in habitats with contrasting predation regimes and population densities result, in part, from local adaptation to different per-capita food levels. We reached this conclusion from a full-factorial laboratory experiment in which fish from two populations were raised to sexual maturity at one of two food levels. We found that fish from Wacissa River, characterised by low-predation/high-density conditions, were larger throughout the experiment and had higher survival to maturity, especially, but not only, at the lower food level, than fish from Trout Pond, a high-predation/low-density population. The increased survival of WR fish was, at least partly, mediated through a larger size at birth. Of the traits measured in our study, only the age at maturity showed no difference between populations. Experimentally reduced per-capita food availability led to smaller body sizes, slower growth rates, delayed maturation, and decreased survival. Size at birth proved a key success factor also in relation to food: a larger size at birth enabled fish to mature relatively early even when they were kept at the lower food level. Lastly, we found that size at birth and juvenile growth were identical in males and females, but that females matured earlier and hence smaller.

Some of these results have been found in earlier studies. The larger size at birth of WR offspring has been reported repeatedly from both field collections (Leips et al. 2009; Leips and Travis 1999; Schrader and Travis 2005; Schrader and Travis 2012) and experimental studies in common gardens (Hale and Travis 2015; Leips et al. 2000; Schrader and Travis 2008). In this experiment, age at maturity did not differ, on average, between fish from TP and those from WR, a result that has been found before (Hale and Travis 2015). Also the smaller size at maturity of fish from TP is consistent with earlier studies (Hale and Travis 2015; Leips et al. 2000), as is the older age at maturation and thus larger size of males compared to females (Hale and Travis 2015). Field data using otolith ring counts as indicators of age also suggest that females mature before males (J. Travis, unpublished data).

Maternal effects, manifested through variation among individual females in the growth and development of their offspring, diminished gradually during adolescence and were no longer detectable by the time that fish matured. This is similar to patterns in other studies of fish species in which maternal effects were important in early life but became less so with offspring age (Heath and Blouw 1998; Venney et al. 2020). This was in contrast to the longer-lasting effects of the differential sizes at birth between females from the two populations. In the comparison between the populations, the smaller initial size of TP offspring carried through to a smaller size at maturity, even though the actual somatic growth rates of immature fish did not differ between the populations. This result illustrates the importance of distinguishing variation in maternal effects within populations from those that might be observed between populations.

The maternal effect within populations, manifested through size at birth, was important in a very particular manner. At low food levels, individuals that were larger at birth matured earlier. This was a substantial effect: an individual that was smaller than 6.5 mm at birth matured ~16 days later than one that was larger than 7.0 mm at birth (see Fig. 5d). This relationship was weaker at high food levels (difference of ~10 days). This result reinforces the conclusion drawn by Leips et al. (2013), who found that larger size at birth was inversely correlated with age at maturity, especially when fish were raised in competitive conditions. The pattern suggests that the larger size at birth in WR fish is an adaptation to low per-capita food levels produced through some combination of lower primary productivity and higher population density.

The most direct evidence that density-dependent selection has been driven at least in part by different levels of per-capita food availability was the stark difference in survival between TP and WR fish at the lower food level. There was a 75% mortality rate for TP fish at low food, compared with a 15% mortality rate for WR fish. At the higher food level, mortality rates were more similar between populations, 15% for TP fish and 0% for WR fish. This interpretation, that WR fish are adaptively more suited to lower food levels, is bolstered by the contributions of the individual cells to the Chi-square score, with the number dead in TP at low food being the major component. Survival was positively associated with size at birth, suggesting that the reduced survival of TP fish resulted at least partially from their smaller average size at birth. From previous work we know that fish from these populations show a trade-off in offspring size and offspring number, with TP females having more, but smaller, offspring than WR females (Leips et al. 2000). Taken together, this suggests that, in TP fish, the production of many small offspring, rather than fewer large ones, pays off because the higher primary productivity of their habitat allows also small fish to survive, while the larger number of offspring ensures that at least some offspring survive to maturity despite the high predation risk.

These results do not preclude other agents of selection from operating differently in the contrasting density conditions in WR and TP. Social stress is important in TP fish (Leatherbury and Travis 2019) but we do not know if WR fish are less sensitive to this source of stress. We also have no information on pathogen incidence and virulence in each population. However, demonstrating that selection at higher density confers a greater ability to cope with low food levels is a step forward in moving the study of density-dependent selection toward a more biological and less phenomenological focus (cf. Engen et al. 2020).

## Acknowledgements

The authors thank Joanna Parrino for conducting the laboratory experiments. This research was funded by a Swiss National Science Early Postdoc Mobility Fellowship (project no. P2EZP3_181775) to A.F.

## Author contributions

J.T. and J.L. conceived the study. J.L. conducted the experiment. A.F. analysed the data, with input from J.T. A.F. and J.T. drafted the manuscript. All authors contributed to subsequent revisions.

## Data Availability Statement

The complete dataset will be available at the Dryad Digital Repository pending manuscript acceptance.

